# Changes in social position predict survival in bottlenose dolphins

**DOI:** 10.1101/2022.08.25.505273

**Authors:** Robert William Rankin, Vivienne Jilla Foroughirad, Ewa Beata Krzyszczyk, Céline H Frère, Janet Mann

**Affiliations:** Murdoch University; Georgetown University; Bangor University; University of Queensland

## Abstract

Social bonds and social structure are important features of animal systems that impact individual fitness. Few studies have examined how temporal dynamics in individual social bonds predict fitness outcomes. This is critical to understand given the high variation in types of social structures and strategies within populations of social mammals, both across time and among individuals. If individually-differentiated social bonds are important, it might be the *change* in bonds which has fitness consequences, not the absolute number of bonds nor their strength. We investigated how network dynamics predict survival in a wild population of bottlenose dolphins using a 35-year longitudinal study. In particular, we were interested in two sex-specific measures of the “widowhood effect”, as well as a more general investigation into the predictability of mortality from changes in higher-order social network metrics. We used two inferential frameworks to provide complimentary evidence for or against hypotheses; namely, a gradient-boosting predictivist approach with relative importance measures, and an inclusion probabilities approach based on stability-selection. We found evidence of a widowhood effect among males but not females. Surprisingly, the most robust predictor of survival was closeness-centrality, whereby the loss of closeness centrality preceded death. This finding, that absolute network position may not be as relevant to survival as changes in network status, is consistent with a large social-psychological literature in humans on the impacts of loss of social capital, dissolution of close friendships, and loss through death of partners (widowhood effect). This study highlights the critical nature of social connections and how its disruption can be a matter of life and death.

## 2 Introduction

In many animal populations, social bonds and social structure are considered to be important mediators of individual fitness and key drivers of evolution. For example, among primates and delphinids, social pressures may be influential in the evolution of large brains [Connor, 2007, Boddy et al., 2012] and complex cognitive abilities [such as tool-use and novel problem-solving; Jolly, 1966, Borrego and Gaines, 2016, Benson-Amram et al., 2016]. Early research on social bonds focused on “non-random” animal associations [e.g., Parsons et al., 2009], where some relationships are individually-differentiated [Lukas and Clutton-Brock, 2018] and transitive [Grosenick et al., 2007], not unlike humans. More recently, multiple studies in mammals and birds have demonstrated how individual variation in sociality contributes to fitness [McDonald, 2007, Frere et al., 2010,?, Mann et al., 2012] and may be heritable [Brent et al., 2013, Gilby et al., 2013, Fowler et al., 2009, Brent, 2015], thus providing early hints about the selection mechanisms that may underlie the evolution of complex sociality and individually-differentiated bonds.

Most of the existing research on social bonds and fitness have used static measures, such as dominance rank, bond strength, or the number of bonds, and their ability to predict changes in fitness-proxies, such as offspring survival, mating success, or longevity. While some individual social traits are relatively stable [e.g., female dominance rank in savanna baboons; Silk et al., 2010], many aspects of animals’ sociality are dynamic and change throughout development [Turner et al., 2017]. Some changes are part of the natural course of maturation (e.g., the expansion or collapse of one’s ego-network) or may be related to important life history events (e.g., changes in reproductive status, migration). A decline in the number of valued social partners or change in one’s social position is also predicted to have deleterious effects [Knight and Mehta, 2017], perhaps through physiological stress resulting from the loss in social status [Creel et al., 2013]. However, no study of a wild animal population has linked such changes to individual mortality events, especially in societies with no clear dominance hierarchy. Here, we examine static and dynamic changes in sociality and social network metrics in a wild population of bottlenose dolphins with the prediction that *changes* (decline) in one’s higher-order social metrics (such as network position and indirect connections) are linked to future mortality events, our proxy for fitness.

The causal direction between changes in sociality, physiology and health, and mortality are difficult to determine from observational data. Much of the literature suggests that fitness benefits are associated with strong social support, although the direction of effects is not known [social buffering hypothesis; Wittig et al., 2016, Silk et al., 2010]. Some researchers have noted incidences where terminal health conditions can lead to large changes in one’s social environment [Miketa et al., 2017], which is a causal direction opposite from our interest in this study. Nonetheless, we suggest that there is likely to be a positive feedback effect, such that an erosion in one’s social connectivity precedes and is predictive of mortality, even if not directly causal *per se*. We formulate this idea within two inferential frameworks, one that is a general proof-of-concept of the predictive importance of changes in sociality (irrespective of the causal direction), and the second which posits specific metrics-as-hypotheses, and which provides evidence in favor or against our hypotheses.

In the case of specific hypotheses, we were interested in whether mortality was influenced by sex-specific social processes, using data from a well-surveyed population of bottlenose dolphins *Tursiops aduncus* in Shark Bay, Western Australia. For the males, who have long-term social alliances [Connor and Krützen, 2015], our conjecture was that there would be a “widowhood effect” [Christakis and Iwashyna, 2003, Elwert and Christakis, 2008, Lillard and Waite, 1995] among closely-bonded male-male pairs (e.g., if my top associate dies in year *t*, then I am at higher risk of dying in *t* + 1). For females, in contrast, who socialize with kin and within specialized foraging sub-communities [Frere et al., 2010, Mann et al., 2012], our conjecture was that females would manifest a similar sort of social-widowhood effect, but driven by the loss of calves and older daughters, rather than social-alliance members, reasoning that daughters serve as top associates and anchors to the broader network. Thirdly, we were also interested in a variety of commonly used social-network metrics and individual-level metrics, building off of earlier work about the importance of social-metrics and fitness-surrogates [Stanton and Mann, 2012, Frere et al., 2010].

However, evaluating hypotheses about social processes using social metrics derived from observational data is fraught with difficulties. The conventional correlation methods and significance-testing frameworks that ecologists use for quantitative support for or against hypotheses have short-comings when used for social-metrics-as-hypotheses. In particular, the Neyman-Pearson hypothesis-testing paradigm suffers low statistical power in social-network studies [Brent et al., 2013]. Furthermore, some social network metrics have multiple operationalizations [e.g., consider the subtle differences among clustering coefficients; Zhang and Horvath, 2005, Opsahl and Panzarasa, 2009] each of which may provide unique information, but whose co-inclusion in statistical analyses results in high-dimensional multi-collinearity. Another concern is that some higher-order metrics (such as centrality, assortativity) may not measure *specific* social processes, but instead are mere network artifacts of lower-level individual metrics [e.g., strength and degree Rankin et al., 2016] or missing data [Firth Josh A. et al., 2017]. This makes the science of social-metrics and fitness-surrogates at risk of over-interpretation: seemingly network-level measures may instead be driven by less-interesting individual-level processes [Firth Josh A. et al., 2017], such as heterogeneity in gregariousness. However, lower-order social metrics, i.e., individual social phenotypes, may have more straight-forward inheritance mechanisms, unlike the network-level indirect connections whose fitness-consequences and inheritance are more controversial [Brent et al., 2013, Brent, 2015].

To address some of these issues, we sought a promising hypothesis-focused framework called stability-selection [Bach, 2008, Meinshausen and Bühlmann, 2010]. It is related to a class of high-dimensional inference methods from the machine-learning community that do well in the face of multi-collinearity and high-dimensionality [Dezeure et al., 2015]. We use this framework to derive “inclusion probablities” that quantify our degree-of-belief in the importance of certain metrics in driving survival.

This inferential framework, however, cannot ameliorate some of the inherent problems of measuring social processes; in particular, many social metrics may not actually measure what they are thought to measure [Franz and Alberts, 2015, Rankin et al., 2016, Firth Josh A. et al., 2017]. This fundamental insecurity casts doubt on the use of *any* hypothesis-driven study of animal sociality using observational data. In contrast to the hypothesis-based inferential frameworks, the *predictivist* framework [Forster and Sober, 1994] side-steps these issues of (over)interpreting metric-as-hypotheses. We therefore propose a predictivist method as a complimentary analysis, in that tests the predictive ability of a model and outcome. It asks the more conservative question: can we predict whether a dolphin lives or dies? This focus on *observables*, and optimization for high-accuracy in prediction, does not demand that there exists any interpretable biological phenomena underlying our social metrics. Rather, predictive performance merely demonstrates that there exists some aspect about social metrics that has *information* about fitness, without yielding rigorous evidence in favor of or against specific hypotheses [Lo et al., 2015]. Like other predictivist tools (e.g., the AIC), this framework can merely provide “relative importance” rankings [Galipaud et al., 2014, Giam and Olden, 2016]. In the following study, we use these two inferential frameworks to garner support for higher-order dynamic social metrics and their effect on mortality, including a high-dimensional hypothesis-focused framework based on stability selection, and a predictivist framework based on gradient-boosting. The two frameworks provide different types of inferences [Shibata, 1986a, Poole, 1987, Yang, 2005, Wit et al., 2012, Vrieze, 2012], which we consider complimentary evidence-based approaches, and which together help mitigate some of the technical and conceptual difficulties of testing hypotheses about sociality using observational data.

## 3 Methods

### 3.1 Field Study Design

The study used 29-years of observations of a resident population of bottlenose dolphins in the eastern gulf of Shark Bay, Western Australia [Mann et al., 2012]. Individual dolphins were identified and tracked over the course of the study (1989 to 2017) through visual recognition (photo ID) of dorsal fin patterns (notches, scars, pigmentation). Associations were estimated from opportunistically encountered groups during boat-based surveys, using a 10-meter chain rule to cluster individuals into groups [Smolker et al., 1992b]. Boat surveys consisted of 2-4 observers on 1-2 boats, with the objective of trying to encounter every cataloged dolphin at least once per season.

### 3.2 Data Preparation

The data included: time-series of individual identifications and dyadic associations during group-encounters; individuals’ mortality status (based on a rules-based assessment of uncertainty, described below); individuals’ estimated age; individuals’ sex; and individuals’ matrilines (clustered into groups that we call haplotypes).

For age estimates, 39.5% of the population had robust birth information. Such individuals were sighted as young calves or as sub-adults. For all other individuals, we estimated age based on physical size and/or skin speckling [Krzyszczyk and Mann, 2012]. Females with no age information who were first sighted with a calf were assigned a birth-date 12 years before the estimated birth year of their first observed calf.

Sex was determined according to: i) genetic data; ii) sightings of reproductive anatomy; and iii) physical proximity between calves and mothers (i.e., females). Individual matrilines were clustered and codified by (inferred) mtDNA haplotype groups, based on generational observations of mother-calf pairs and genetic samples (hereafter, simply referred called “haplotypes”).

For the analyses’ response variable, individuals were assigned a dichotomous mortality status: (*alive* or *dead*) during two-year time-windows. We also assign an uncertainty grade to each mortality status: >95% probability of being alive, 72% probably alive, 50% alive or dead (e.g., unknown), 72%probably dead, and >95% dead. 13% of individuals had *final* mortality status that were less than 95% certain of being alive or dead. Uncertainty classes were assigned based on multiple lines of evidence, such as disappearance following observations of acute ill-health (e.g., shark wounds, emaciation), or sightings of an individual’s orphaned progeny (e.g., for mother-calf pairs). Finally, individuals who were absent from the sighting-catalog for at least three years were coded as “dead” or “probably dead” using a rules-based procedure: individuals who were absent more than the longest gap in their individuals’ sighting, by a multiple of three, were assigned a death status. For example, if an individual had a maximum observed-absence of three consecutive years, we waited 9 years to assign “dead” with 95% certainty. A similar logic followed “probably alive/dead”. Otherwise, individuals were coded as unknown (50% probability of being alive or dead). Using this method, no individuals have turned up ‘alive’ after having been assigned a high-certainty of death.

The mortality-certainty categories were incorporated into the analyses via weighted logistic regression, whereby the two outcomes (alive, dead) had a combined weight of 1, and their respective weights reflected our certainty about an individual’s mortality status: e.g., if an animal had a probability of being dead=0.72, then one “death” observation incremented the log-likelihood with a weight of 0.72, while one complimentary “alive” observation incremented the log-likelihood with weight 0.28.

Finally, we truncated the sample of individuals to minimize several sampling biases. The truncated dataset included individuals who were: i) alive post-weaning; ii) sighted at least 20 times during the entire study; iii) sighted in at least 5 consecutive years. Animals not meeting these criteria could potentially have caused a negative-bias to the survival estimates, given that their mortality status would be highly confounded with geographic movement or changes in observational effort. The truncation resulted in 465 unique individuals and 249 death-events. The “class imbalance” between observations of death versus observations of being alive motivated us to re-weight the two classes for the predictivist model (below).

### 3.3 Hypotheses and Social Metrics

In the statistical analyses, we used a two-year moving-window to summarize data. For social metrics, this meant that social networks were constructed using Half-Weight (HW) associations [Cairns and Schwager, 1987] over two-year windows, per year. For instance, an individuals’ metric-value at year *t* was calculated based on information from years *t-1* to *t*. If an individual was missing for both years within a window, the data was set to “missing” during that interval. An animal’s age within a window was set as the mid-point of *t-1* and *t*. For mortality status, individuals were assigned a value of “dead” for the last window in which they were observed (subject to the uncertainty bins above). In other words, if a death-event occurred anytime during *t* or after *t*, it was assigned to the window *t-1* to *t*: this ensured that social information from lag 1 to the present *t* were being used to predict the probability of death at anytime during or after *t*. ^1^

### 3.4 Metrics

The covariates included six types of metrics: individual-level covariates, global social-network metrics, sex-based homophily social-network metrics, change in social metrics, and metrics designed to address specific hypotheses about social processes.

#### Individual-level metrics

Individual-level covariates included features like age, sex, and “haplotypes” (matrilineal and mitochondrial clusters and). In this category, we also included social metrics that were not network-based but were measured at the individual-level, such as: “group-size” (average group size per individuals’ encounter); “p(alone)” (the proportion of encounters when an individual was alone), strength (sum of weighted associations), degree (sum of unweighted associations), and disparity [individuals’ variance of network-weights Barthélemy et al., 2005].

#### Global social-network metrics

This class of metrics included higher-order measures of indirect connectivity and individuals’ positions within networks, such as: weighted centralities [betweenness, closeness and eigenvector; Freeman, 1979, Butts, 2008, Opsahl et al., 2010], weighted clustering [Opsahl and Panzarasa, 2009], affinity [nearest-neighbor degree/strength; Whitehead, 2008, Kasper and Voelkl, 2009], and assortativity (affinity divided by degree/strength).

#### Homophily network metrics

For the above network covariates, their values were calculated based on the entire social-network (i.e., a network including all individuals, regardless of sex). We also calculated sex-homophily versions, i.e., *c*_*i*_ = ∫_*τ*_ closeness(𝒢_*τ*_)_*i*_*dτ*, where 𝒢_*τ*_ was a weighted graph whose edges *w*_*ij*_ between opposite sex pairs *i* and *j* were down-weighted by a scalar *τ* ∈ (0, 1]. The motivation for the sex-homophily metrics was to emphasize the indirect connections among same-sex nodes, while avoiding the computation problems that arise from disjointed sex-exclusive graphs. In cases where the homophily and non-homophily metrics were highly correlated (> 0.9), we used predictive performance to judge which covariate to discard (i.e., a change in cross-validation log-likelihood). Covariates that were retained are shown in Figure 1.

#### Change in social metrics

All the aforementioned social metrics were calculated as the absolute values per 2-year time-window. We also computed dynamic versions: the difference between each time-windows’ absolute value versus an individuals’ long-term mean (a.k.a., deviations from the mean). Such deviations were meant to approximate the difference in values between consecutive time-windows. Unfortunately, the differences could not be calculated because of gaps in some individuals’ sighting record (e.g., due to temporary migration or differences in observational effort, hereafter simply referred to as “statistical temporary emigration”). The correlation between the deviations versions and the time-differencing versions was high ≫ 0.7, and thus we accept our dynamic method as an approximation of the time-differencing approach.

#### Standardization

For the social-network metrics, a final consideration was how to normalize for artefacts due to year-to-year changes in networks’ degree [Franz and Alberts, 2015] or other year-effects. We tried two types of metric normalization: z-score normalization and the median absolute-deviation. A cross-validation exercise suggested that the raw metrics (no normalization) had the highest predictive power.

#### Metrics as hypotheses

The final category of covariate pertained to our specific hypotheses about the importance of social structure and social processes on bottlenose dolphin mortality. The first covariate operationalized a “widowhood effect”: that long-term male-male alliances have evolved with fitness benefits, such that when an individual is “widowed” and unlikely to re-establish a close bond, his probability of dying increases. The metric was calculated as the sum of an individual’s associates who died during the previous time-window weighted by the associates’ HW^2^ values (and logged). The metric is high when an individuals’ top-associate dies (i.e., they are widowed), or if many low-ranking associates die.

Another three covariates pertained to our hypothesis that adult females’ social-standing and connectedness to the female community was maintained by her calving-state and matriline. The first of three covariats was “Calf death”: a binary indicator whether a mother lost a calf during a time-window. “Weaned daughter death” was a binary covariate which scored whether a mother had lost any weaned female offspring during a time-window (excluding cases where the mother and offspring may have disappeared around the same time). “Surviving weaned daughter” was a binary covariate which scored whether a mother had any surviving weaned females. Changes in these states were hypothesized to increase a mother’s mortality through social isolation, like a widowhood effect betweeb mothers and daughers/calves.

Note, while these metrics tried to target causal social processes, whereby the loss of close-associates has negative social consequences on survivors, there was some inevitable confounding with possible injurious events which may have killed both mother and offspring at different times, rather than one’s death being due to a social process. For example, a shark attack which immediately kills a calf and injures a mother, such that the mother dies at a latter date. All widowhood metrics potentially had this type of confounding.

In total, we used 57 covariates; many with high correlations among each other. For example, the maximum correlation between all covariates was 0.84, thus necessitating the following shrinkage and regularization methods.

### 3.5 Predictivist Statistical Inference

For the predictivist approach, we compared three types of weighted-logistic regression techniques: component-wise gradient boosting [Bühlmann and Yu, 2003], support vector machines [Cortes and Vapnik, 1995], and deep neural networks with entity embeddings for categorical variables [Chollet et al., 2015, Abadi et al., 2015]. Based on cross-validation performance, we decided to use component-wise gradient boosting.

Component-wise gradient boosting is a type of ensemble-building and *𝓁*_1_-regularization technique [Bühlmann and Yu, 2003, Efron et al., 2004, Mayr et al., 2014]. A boosted ensemble consists of thousands of weak “base-learners”. We used three types of learners: i) penalized least-squares learners (PLS) for all categorical variables (sex, haplotype, time-as-a-factor, calf deaths, weaned daughter deaths, surviving weaned daughters); ii) PLS learners for interactions between the aforementioned covariates and age; and iii) conditional inference trees [Hothorn et al., 2006] with an interaction depth of 2 (set by cross-validation) for all covariates (except year-as-a-factor, implying that the effect of other covariates were stationary in time).

Component-wise gradient boosting has a few theoretical and empirical properties which are relevant for studying social metrics. First, *𝓁*_1_-regularizers (like the lasso and component-wise boosting) provide a principled means of automatic variable selection, motivated by maximizing predictive performance: base-learners that contribute to predictive accuracy have more weight; variables that do not are shrunk towards zero. Such shrinkage is a way of negotiating the bias-variance trade-off to prevent over-fitting [Schmid et al., 2010]. Furthermore, *𝓁*_1_-regularizers are capable of handling high-dimensionality [Schmid et al., 2010], and are more robust to multi-collinearity, compared to alternative model-averaging methods [Strobl et al., 2008, Cade, 2015][but they require additional tweaks for inference Bühlmann et al., 2013]. The latter property is particularly important for social-network studies where collinearity among metrics is onerous [Brent et al., 2013].

The high predictive performance of component-wise gradient boosting [Chen and Guestrin, 2016, Guyon et al., 2009] can be understood by understanding its objective function: the expected negative log-likelihood 𝔼[−log(ℒ)] [Bühlmann and Yu, 2003], a.k.a, the Expected Risk.^2^ It is important to understand that minimizing 𝔼 [−log(ℒ)] has theoretical optimality properties for prediction and estimation, but not for hypothesis-driven inference, which is the emphasis of the next section [i.e., such methods have comparatively high Type-I errors; Shao, 1993, 1997].

Through the lens of predictive accuracy, we scored covariates based on their relative variable importance (RVI). We call this “weak inference”, given that the theoretical properties of RVI statistics are poorly known [Galipaud et al., 2014, Giam and Olden, 2016, Gregorutti et al., 2017]. Three RVI statistics were computed: i) the in-sample explained-risk reduction RVI [Elith et al., 2008], which scored covariates based on their contribution to the improvement in the Empirical Risk (the in-sample negative log-likelihood) during gradient descent; ii) an out-of-sample version of *(i)*, which used out-of-sample data to compute each covariates’ contribution to the Risk reduction, averaged over 100 bootstraps; and iii) the drop-out RVI, which dropped whole groups of correlated/related covariates and measured the degradation in predictive performance (change in Expected Risk). The intuition for *(iii)* is the same as the popular Breiman score Breiman [2001]. RVI’s *(i)* and *(ii)* were useful for scoring the degree to which covariates worked together internally to the variable-selection algorithm, whereas RVI *iii* only looked at the unique contribution of a group of covariates which could not be explained by all combinations of other covariates.

The key regularization hyper-parameters were: the number of boosting iterations, the learning-rate, and the interaction depth of conditional-inference trees. We also considered the class-weighting for the imbalanced classes (dead, alive) to be a type of hyper-parameter. These were tuned via a 100-fold bootstrap validation (on average, this left-out 36.6% of the data for validation per bootstrap).

### 3.6 Hypotheses

Once we verified that our suite of social covariates had good predictive power for dolphin mortality, we employed stability-selection [Bach, 2008, Nicolai Meinshausen and Peter Bühlmann, 2010, Shah and Samworth, 2013] for inference about specific metrics-as-hypotheses.

Stability selection was originally conceived as a sub-sampling technique to modify *𝓁*_1_-regularization estimators for the purpose of controlling variable-selection error-rates in high-dimensional settings. It has been used to bound the expected number of false discoveries [Shah and Samworth, 2013, Hofner et al., 2015], or obtain statistical consistency [Bach, 2008], or, in our application, to approximate “inclusion probabilities” [specifically, frequentist approximations of Bayesian posterior inclusion probabilities; Draper, 2010, Murphy, 2012], i.e., *S*_*k*_ ≈ *π*(*β*_*k*_ ≠ 0|**Y**, **X**). Our interpretation of *S*_*k*_ is subjectively (and approximately) Bayesian in the sense that we interpret *S*_*k*_ as being the quantification of our uncertainty about the system. The *S*_*k*_ represent our confirmation functions [Hawthorne, 2011] such that high values (*S*_*k*_ » 0.9) indicate which base-learners are our best current candidates for believing as truly important, conditional on the data and the implicit exponential prior^3^ [Bach, 2008] suggests using a cut-off of *S*_*k*_ 0.9 in order to obtain statistical consistency.^4^

In contrast to the predictivist framework, consistent estimators are not necessarily good at prediction [Shibata, 1986b, Lo et al., 2015]. Furthermore, for stability selection, we could only use linear base-learners, not decision trees, which further has negative consequences on predictive ability.

For stability selection, the universe of base-learners consisted of two types: all covariates in univariate penalized least-squares learners (PLS), and all pairwise-interactions in PLS with a fixed degrees-of-freedom of 1. The latter penalization was necessary to ensure that the learners were “exchangeable” [an assumption of stability selection; Meinshausen and Bühlmann, 2010], and that the component-wise boosting selection mechanism was not biased to multivariate learners (which are inherently more flexible and have a positive selection bias if not penalized).

We calculated inclusion probabilities using the mean operator [Shah and Samworth, 2013] and bootstrap sampling [also mentioned in Shah and Samworth, 2013] instead of the original sub-sampling method of Meinshausen and Bühlmann [2010].

### 3.7 Confounding Factors

#### 3.7.1 Statistical Temporary Migration

Research that use photographic capture-recapture methods to study survival face a major confounding factor called statistical temporary-emigration: when missing animals are mistakenly declared to be dead. One way we attempted to address this issue was by including weights on mortality status based on our assessment of animals being alive or dead.

Another way we tried to address this issue was by conducting an auxiliary analysis. We devised an index which scored covariates based on whether or not they were more linearly associated with known cases of temporary-emigration (over time-durations of 3, 4, 5, 6 and 7 years) vs. linearly associated with death. This was possible by leveraging the length of the time-series to investigate specific cases where an individual was absent for an extended period of time and who later re-entered the dataset (and thus, may have been mistaken for dead, had the time-series been shorter). The linear associations were calculated with simple logistic regression models, whose outputs were the coefficients of association between covariates and outcomes. We then calculated a “mortality premium index”: an index for each covariate based on its comparative degree of association between mortality vs. temporary emigration. A low premium meant that a covariate was comparatively more associated with temporary emigration than death.

#### 3.7.2 Data Leakage

Social metrics may be sensitive to confounding factors like emigration, poor health, or non-random survey effort, if such individuals become less prevalent in the data-set prior to their disappearance/death for reasons unrelated to changes in sociality. The most sensitive metric is degree, because: i) most social-network metrics are sensitive to degree, and ii) in fission-fusion societies, animals with more survey-encounters generally accrue more observed associates, regardless of their actual gregariousness. Therefore, if we observed a large predictive importance to change-in-degree, this may have indicated “data-leakage”, whereby an animal’s absence/death affects the backwards-calculation of social metrics through reduced degree, but not necessarily due to some changed social process.

### 3.8 Age

Age is an importance predictor of mortality. It is likely to have additive or multiplicative effects in combination with other social metrics, but may also be confounding as well. For instance, older animals are more likely to both die and have lost their close associates due to old age. The risk is that we may attribute a causal connection between mortality and a metric that is just a proxy for aging. Ideally, from an ease-of-interpretation perspective, social metrics and aging should impact mortality in independent ways, which may be discernible through three-interaction plots between the log-odds of mortality, age, and a metric of interest.

## 4 Results

### 4.1 Model Assessment

We used the following statistics to assess and compare overall models’ predictive performance: the Expected Risk 𝔼[−log(ℒ)] (approximated with a 100-fold out-of-sample bootstrap validation); the in-sample Area Under the receiver-operator Curve (AUC); the 100-fold out-of-sample AUC (cv-AUC); the in-sample Precision-Recall Curve (PRC); and the out-of-sample PRC (cv-PRC).

Among the three predictivist modeling techniques (SVM, deep neural networks, and component-wise gradient boosting), the SVM had the lowest Expected Risk (0.1971). However, gradient boosting had the higher AUC and PRC statistics, for both in-sample and out-of-sample statistics (AUC 0.943, cv-AUC 0.867, PRC 0.552, cv-PRC 0.3346), and only slightly lower Expected Risk (0.199) as compared to SVMs. Therefore, we used the gradient boosting model for our predictivist inferences. The hyperparameters of the final model were: 948 booting iterations, a learning rate of 0.07, and a tree-depth of 2. To address the imbalanced class-weights, we used a ratio of 1:2.5 for the alive vs. dead outcomes, which was also tuned by cross-validation.

For comparison, the stability-selection model had an in-sample AUC of 0.873, a cv-AUC of 0.834, an in-sample PRC of 0.371, and a cv-PRC of 0.284. This hypothesis-based model had lower predictive performance for two reasons. First, it was not optimized according to a predictivist objective function, but instead sought the most stable covariates for inference about covariates-as-hypotheses. Secondly, it only only included two-way linear interactions, for the sake of interpretability, unlike the predictivist model which included non-linearities and multi-variate interactions.

### 4.2 Variable Importance and Inclusion Probabilities

Among the different relative importance indices (RVI), there was broad agreement about the high importance of age, Δ-closeness centrality (sex-homophily version) and the widowhood effect metric (sex-homophily version).There was some inconsistency about the importance of Δ-p(alone), haplotype and sex. The differences among RVI seemed to be due to whether or not an importance index was based on the *realized* variable-selection behavior, such as the explained-risk reduction RVI [Elith et al., 2008] and the inclusion probabilities (Figures 1 and 3), in contrast to the the drop-out RVI, which looks at the marginal decrease in predictive performance from withholding covariates.

**Figure 1.**
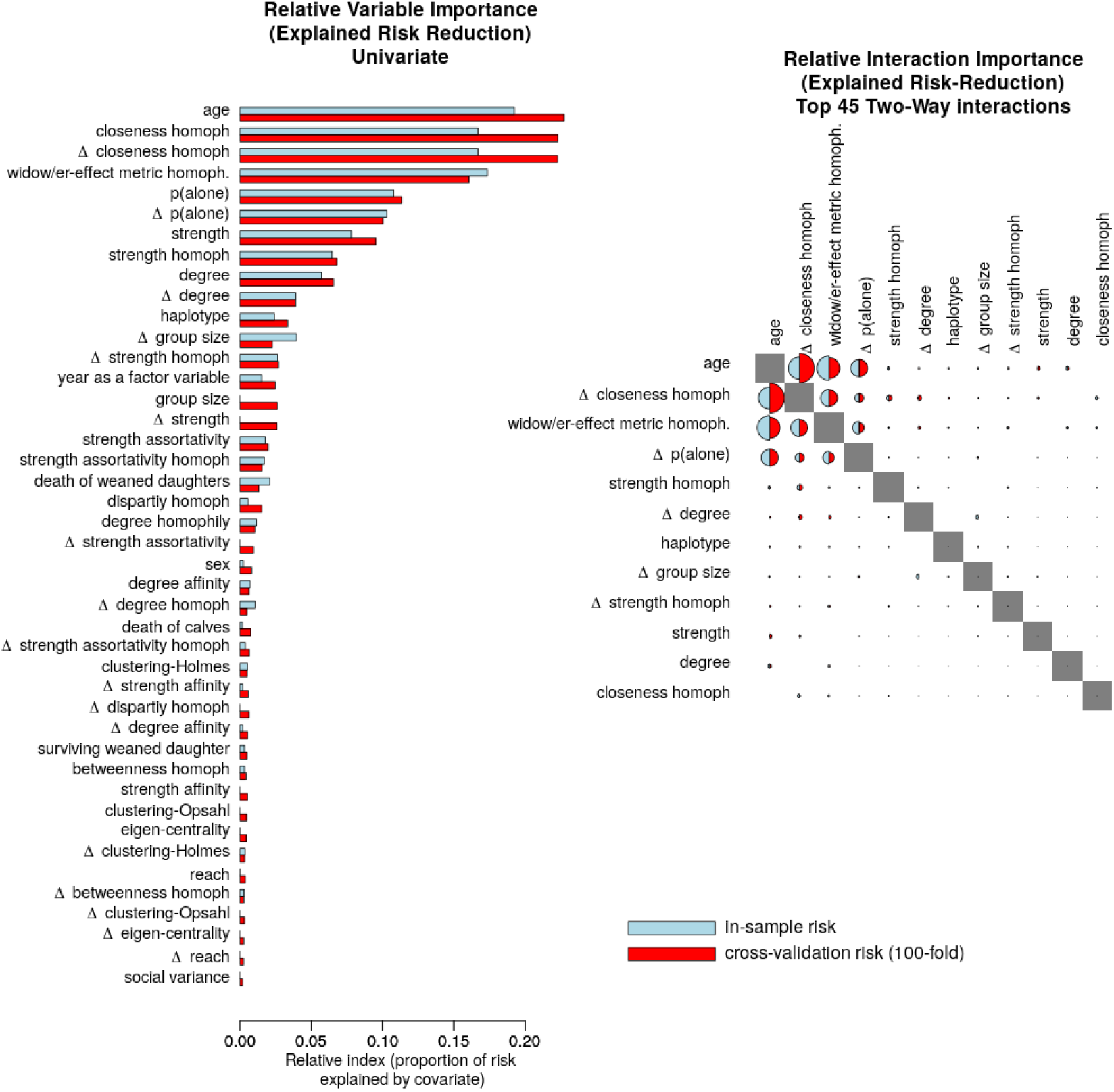
*Left* : Relative variable importance from component-wise boosting, as measured by each covariate’s contribution to the reduction in Risk. *Right* : Relative interaction importance, whereby each circle’s diameter is proportional to the explained risk reduction. *Red* is the index measured on hold-out risk (averaged over 100 bootstraps) and *blue* is the index measured on the “empirical risk” using all the data.

**Figure 2.**
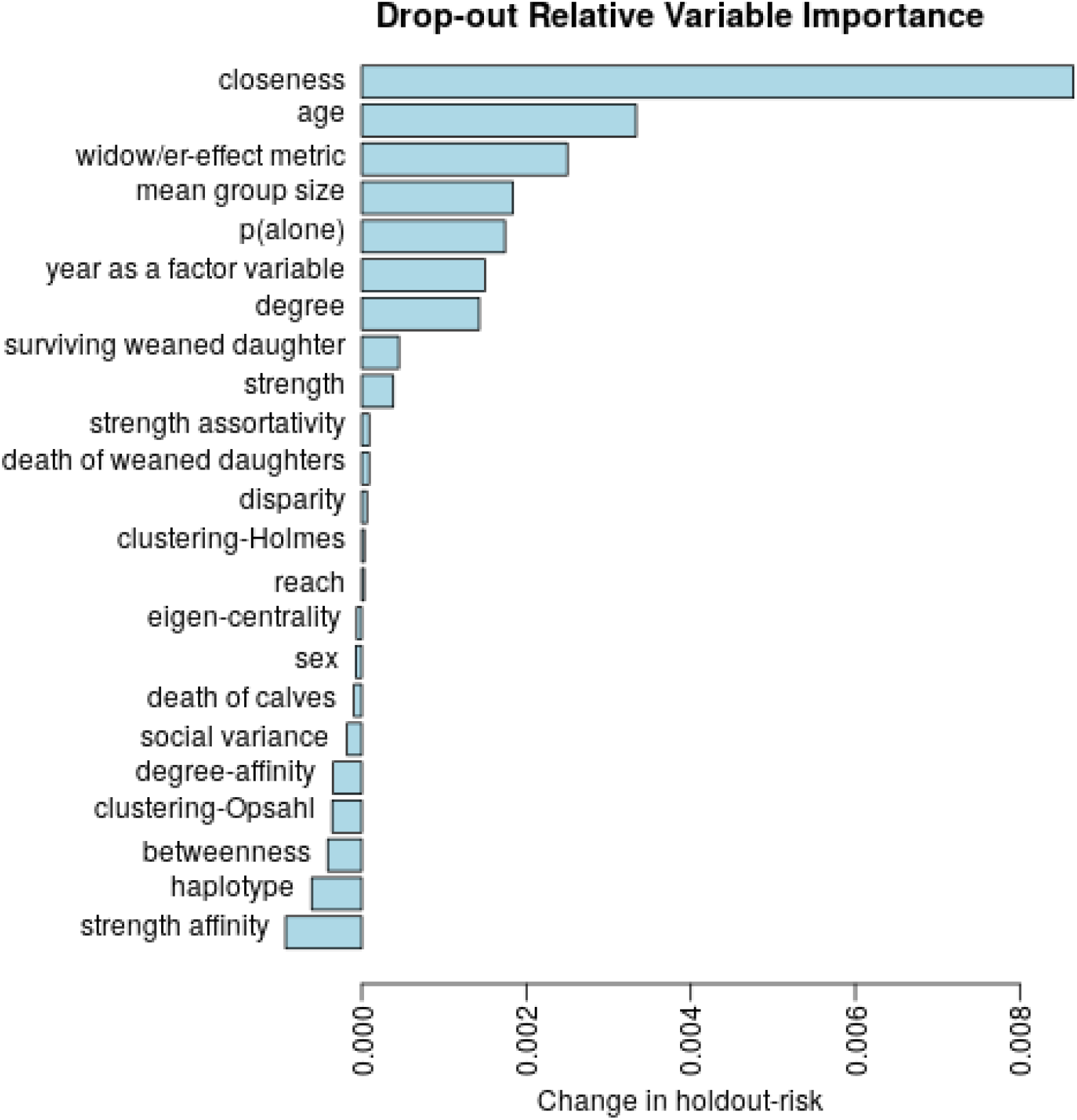
Drop-out relative variable importance, as measured by deleting each covariate group and estimating the change in Expected Risk from the baseline model. Higher positive numbers suggest high importance. Negative numbers suggest that removing the covariate actually improves the predictive importance.

**Figure 3.**
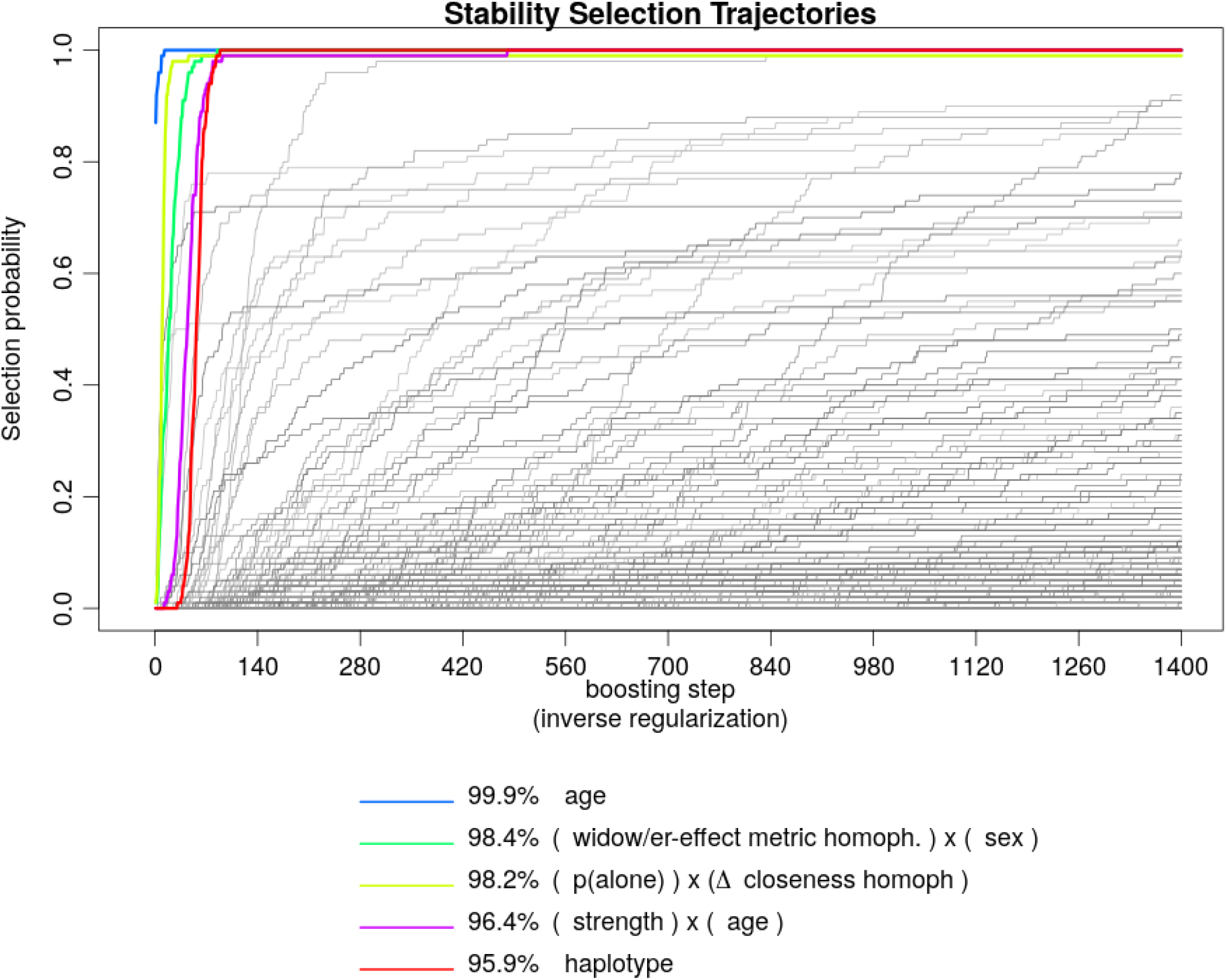
Stability selection pathways and inclusion probabilities of base-learners in component-wise gradient boosting, approximated over 100 bootstraps. Base-learners with lines in the upper-left quadrant have more stable selection overall subsets of the data. Higher inclusion probabilities deserve higher credence.

The explained risk-reduction RVI (Figure 1) gave the highest relative importance to a cluster of six interacting covariates: age, Δ-closeness centrality (sex-homophily version), the widowhood effect metric (sex-homophily version), Δ-p(alone), and, to a lesser extent, strength and Δ-degree. Figure 4 shows an example tri-variate interaction between age, sex and the widowhood effect metric, while Figure 5 shows the interaction between sex, Δ-degree and Δ-closeness centrality. Two more tri-variate interactions are shown in Figures 6 and 7. As expected, the log-odds of dying increased monotonically with age. The widowhood effect was most apparent in males, where there were many more death-events in the high-value region of the metric space (see the orange marks in Figure 4), as opposed to, females for whom this pattern was absent.

**Figure 4.**
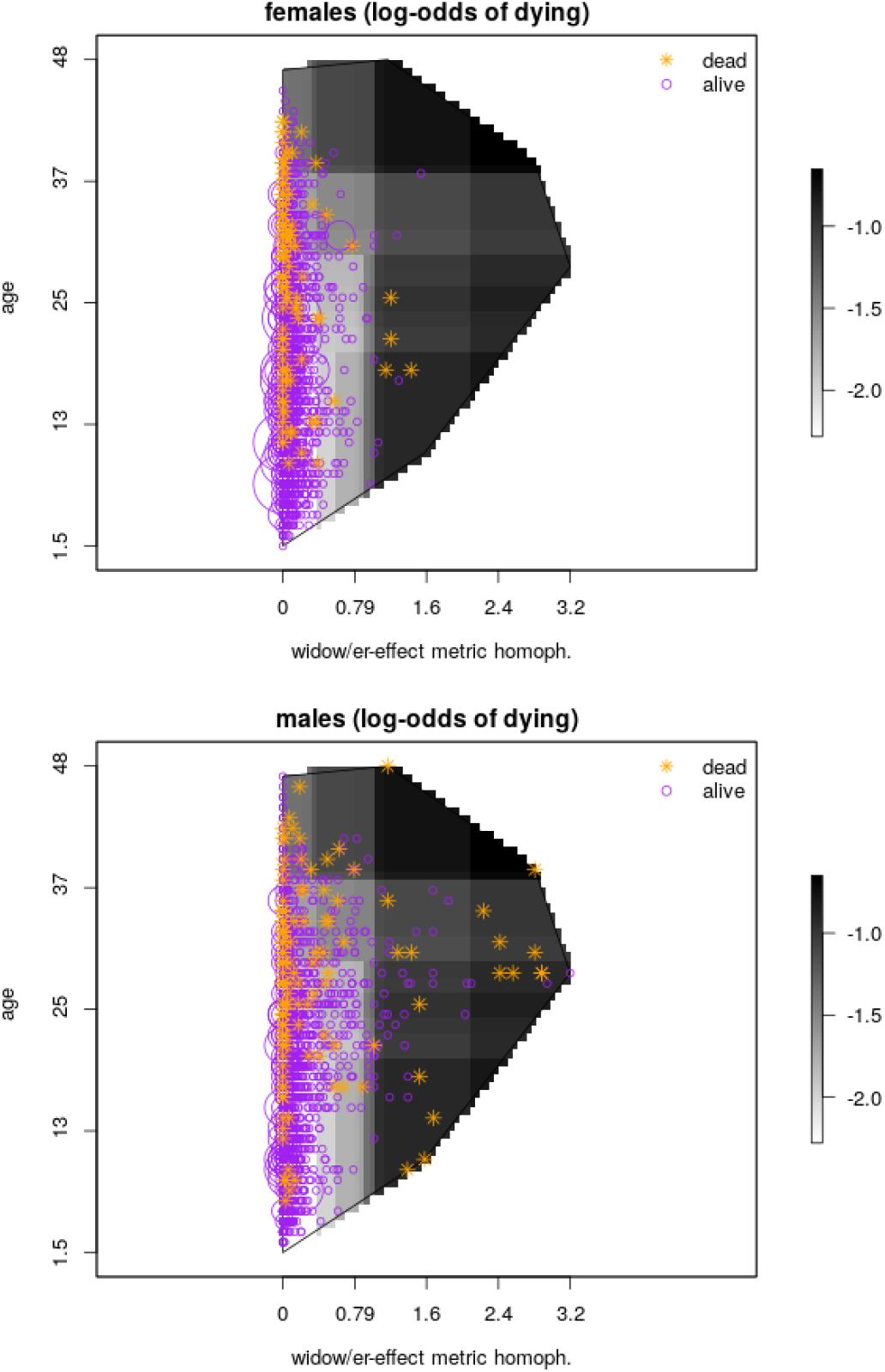
Three-way interaction between the widowhood effect metric (*x-axis*), age (*y-axis*) and sex (*top vs. bottom panels* on the log-odds of dying. Dark colors represent higher estimated log-odds. The overlay of purple and orange dots represent the outcomes (alive and dead, respectively) used to train the models. Note: the boundary of the 2D covariate-space has been clipped to avoid extrapolation into unobserved values of the covariates.

**Figure 5.**
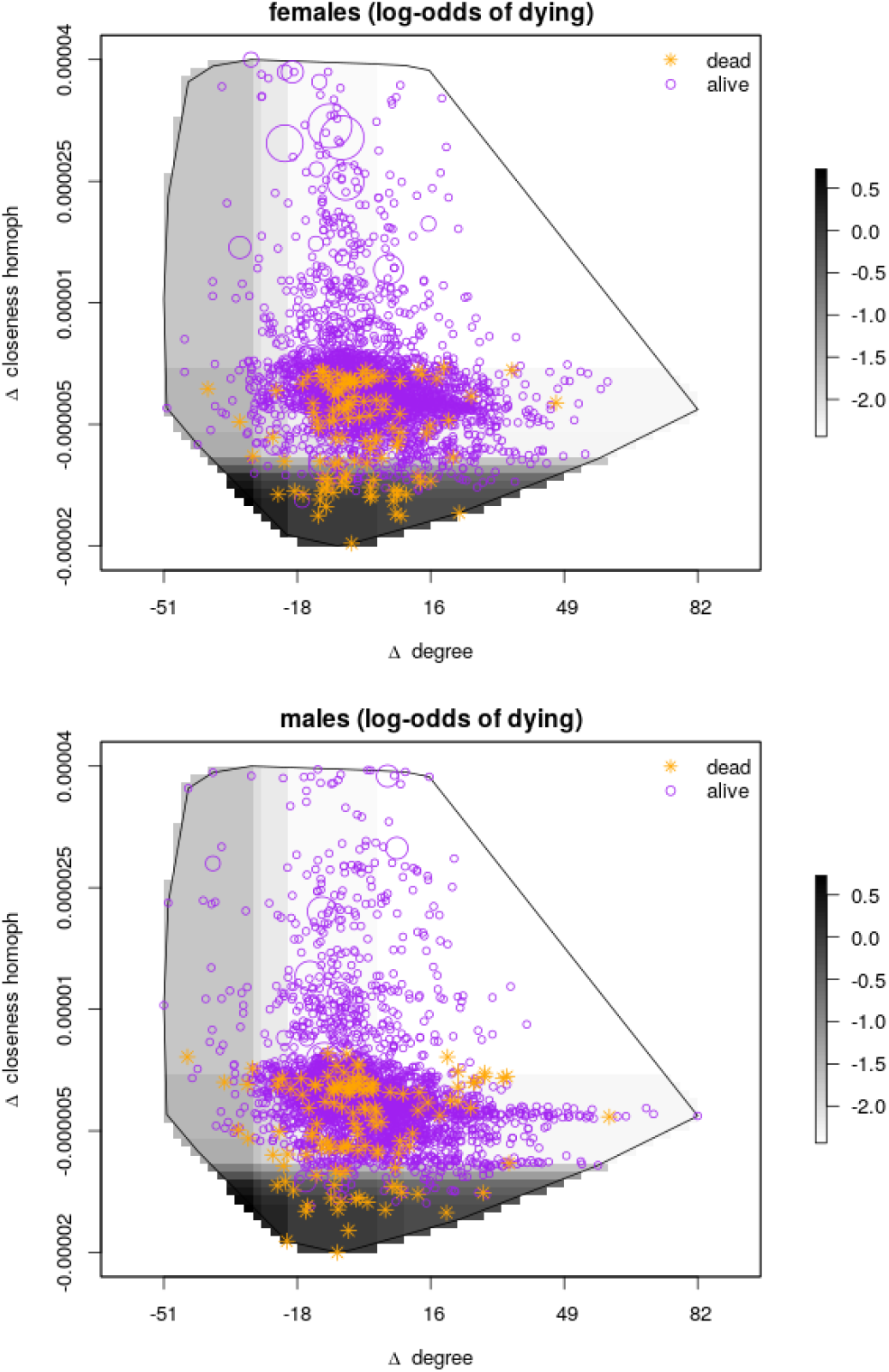
The estimated log-odds of dying according to a three-way interaction between age (*x-axis*), Δ - closeness centrality (*y-axis*) and sex (*top vs. bottom panels*. Dark colors are higher estimated log-odds of dyning. The outcomes (alive and dead) are overlaid as purple and orange dots, respectively. Note: the boundary of the 2D covariate-space has been clipped to avoid extrapolation into unobserved regions of the covariate-space.

**Figure 6.**
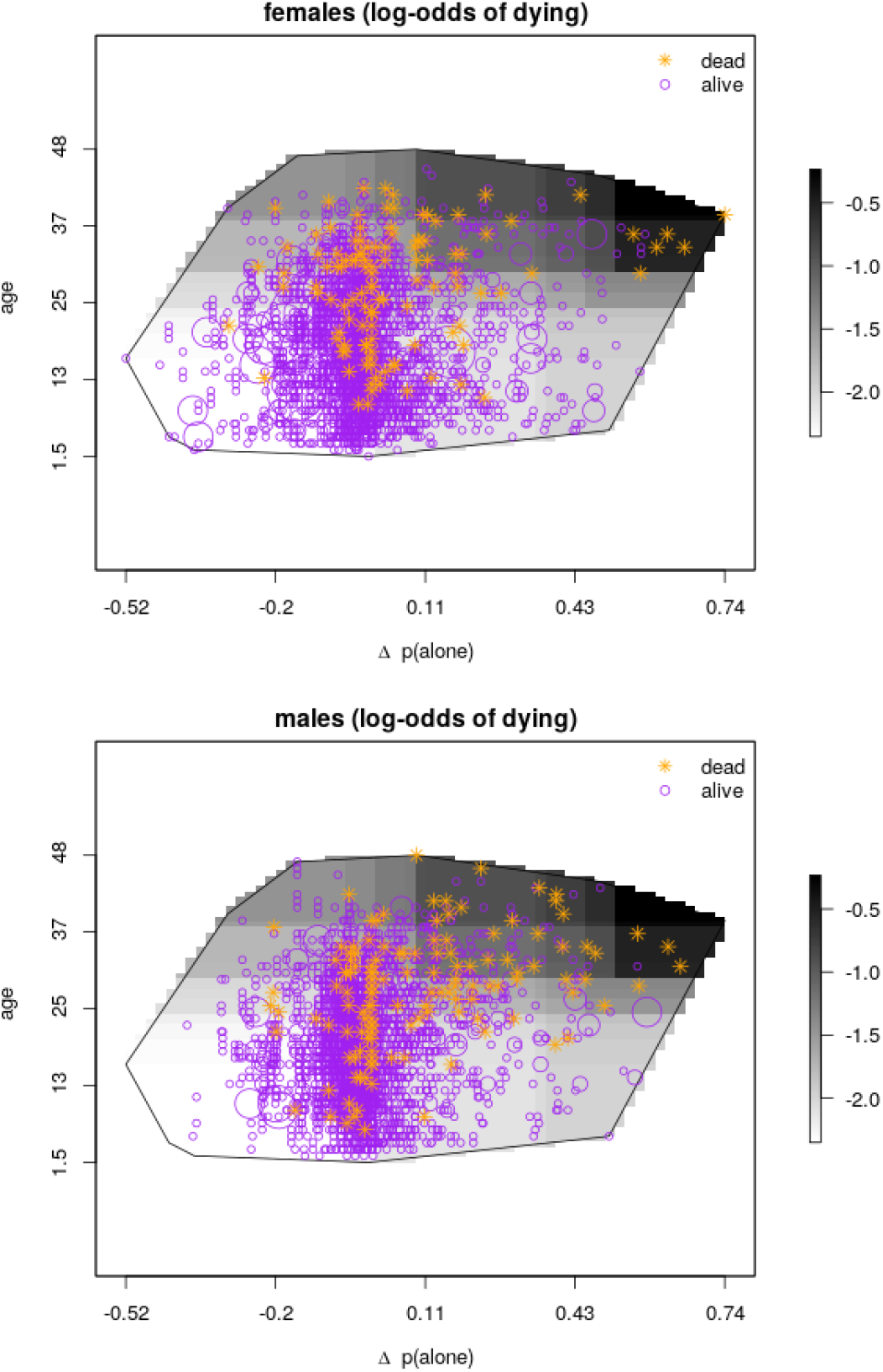
The estimated log-odds of dying according to a three-way interaction between age (*y-axis*), Δ - p(alone) (*x-axis*) and sex (*top vs. bottom panels*. Dark colors are higher estimated log-odds of dyning. The outcomes (alive and dead) are overlaid as purple and orange dots, respectively.

**Figure 7.**
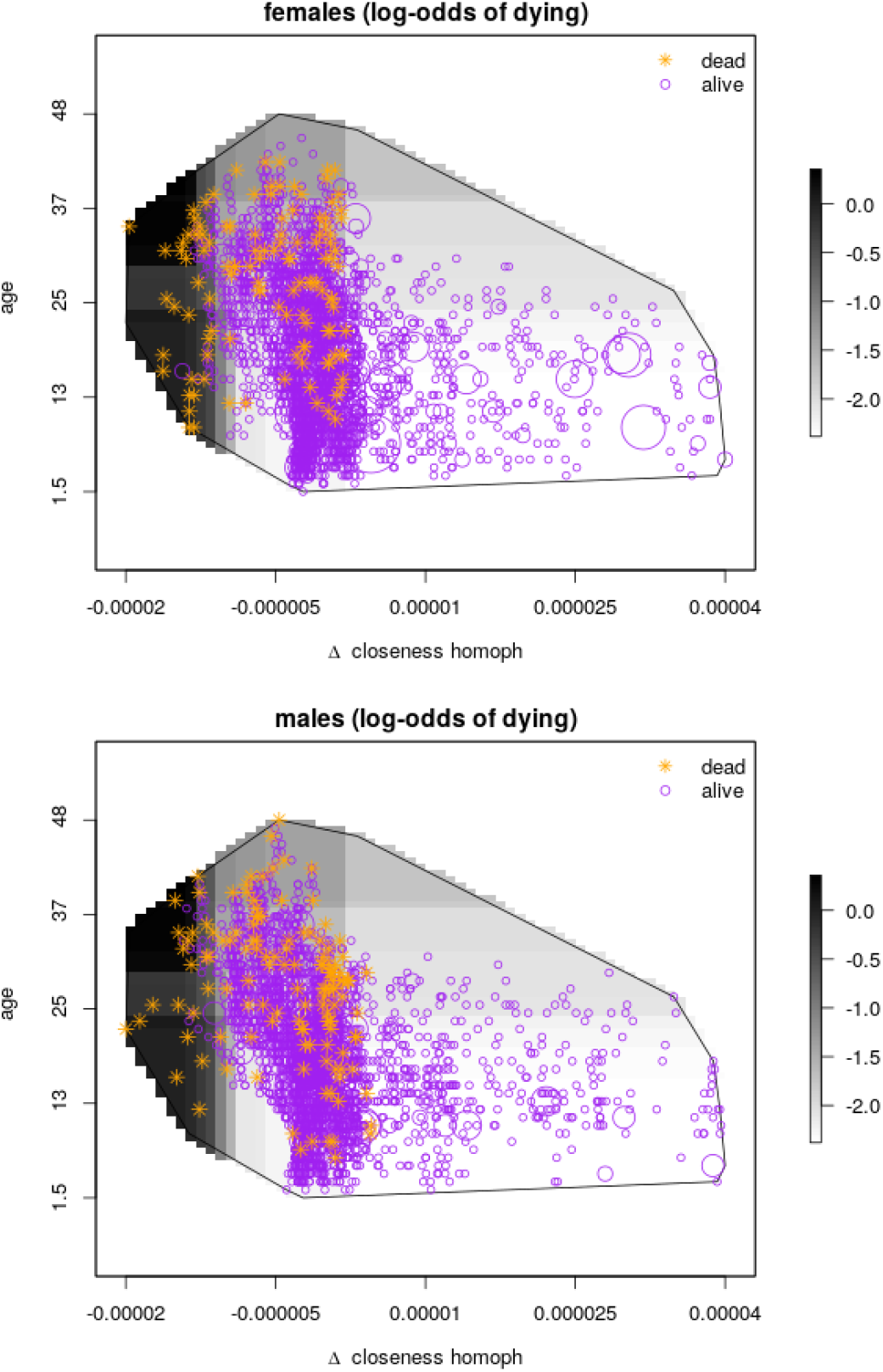
The estimated log-odds of dying according to a three-way interaction between age (*y-axis*), Δ - closeness centrality (*x-axis*) and sex (*top vs. bottom panels* on the estimated log-odds of dying. Dark colors are higher estimated log-odds of dyning. The outcomes (alive and dead) are overlaid as purple and orange dots, respectively.

Δ-closeness centrality (Figure 5) showed the largest differential in the log-odds of dying, and also showed the most dramatic separation between “dead” and “alive” outcomes: almost no death-events occurred above the median Δ-closeness value, whereas deaths were heavily concentrated in the bottom decile of the metric space. The interaction with Δ-degree was almost non-existent (despite the latter’s high RVI score), which has important implications for diagnosing the model assumptions and data-leakage (see below).

According to the dropout-RVI, the closeness centrality metrics had the highest unique importance, both according to its absolute-value metric and its dynamic metric. As a group, the closeness metrics had double the RVI value as compared to the next most important covariate (age). The widowhood effect metric, mean group size, p(alone), year-as-a-factor variable and degree, all demonstrated some unique contribution, albeit with much less relative importance than closeness. Some covariates had negative contributions (such as haplotype and sex), meaning that we could have improved the predictive performance by discarding these covariates altogether.

The approximate inclusion probabilities and their selection trajectories are shown in (Figure 3). The inclusion probabilities for individual covariates were: age >0.999, sex 0.985, (log) widowhood effect metric 0.9853, p(alone) 0.983, Δ-closeness 0.983, and haplotype 0.9596. If we adopt an approximate Bayesian interpretation [Draper, 2010], we should regard these covariates as our best current candidates for the ‘true’ model. Alternatively, if we adopt a frequentist and consistency framework and use the recommended decision rule of keeping covariates whose inclusions frequencies exceeding 0.90 [e.g., Bach, 2008], we would obtain long-run Type-I and Type-II errors that are minimized^5^. A plot of the realized selection trajectories [Nicolai Meinshausen and Peter Bühlmann, 2010] suggest that there was not an obvious cut-off between low- vs high-stability baselearners; we interpret this result as evidence of high model uncertainty. This is undertandable according to the insights from Bühlmann et al. [2013], who notes that the in-sample selection of base-learners is somewhat arbitrary when there is a high degree of multi-collinearity.

### 4.3 Confounding Factors

#### 4.3.1 Statistical Temporary Migration

Regarding the issue of statistical temporary migration, Figure 8 shows the indices of linear association between covariates and mortality as well as statistical temporary emigration. The index “premium” is the degree to which a covariate is more strongly associated with mortality versus migration. The results were consistent with the RVI studies: those covariates with high RVI also had a high mortality premium, and were unambiguously *not* associated with statistical temporary emigration.

**Figure 8.**
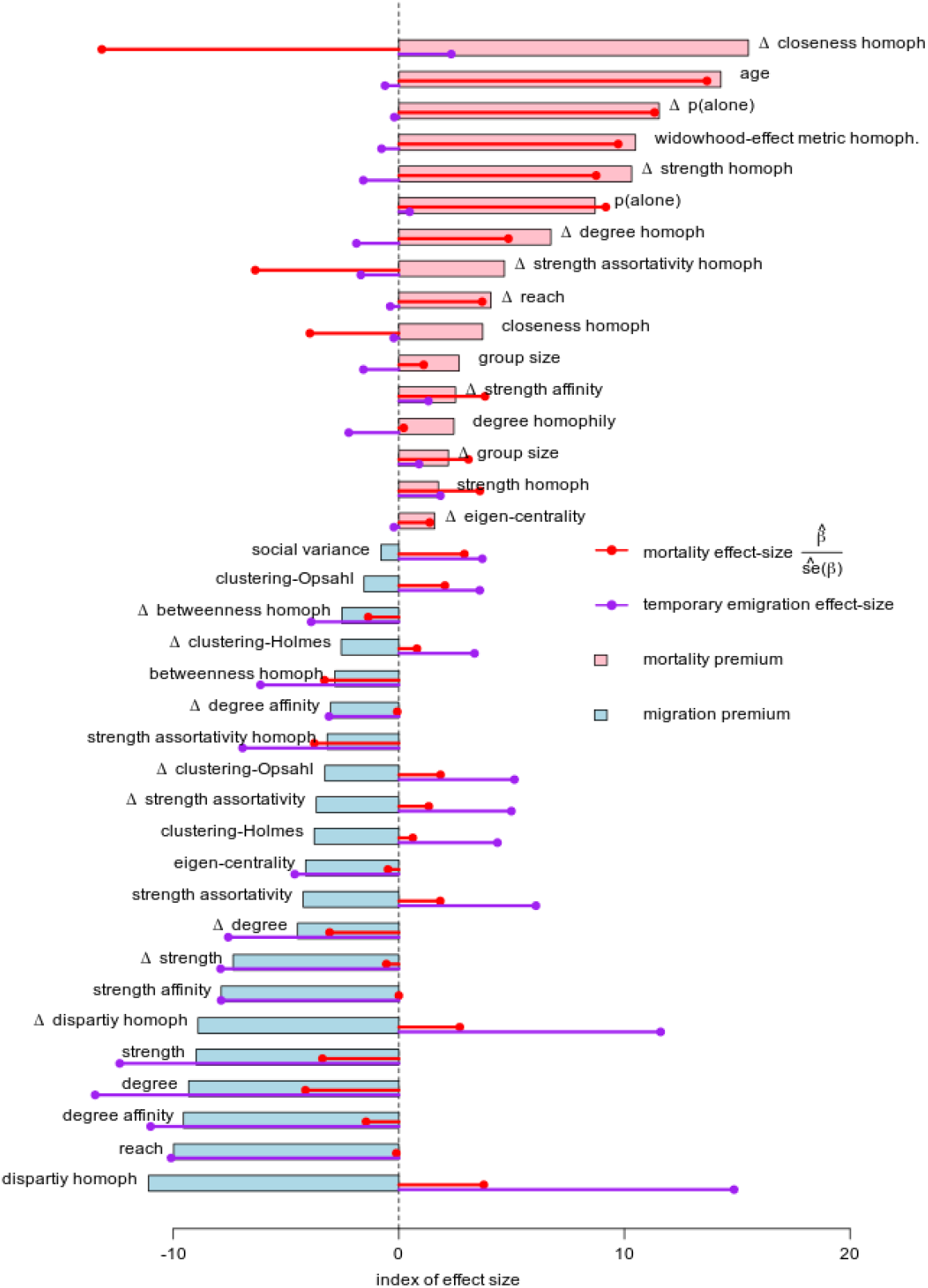
Index of the explanatory ability of covariates (*y-axis*) impacting either mortality (*red-lines*) versus statistical temporary emigration (*blue-lines*). Red and blue lines show the standardized coefficient of association between a covariate and mortality and emigration 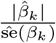. Colored bars show the mortality premium index: the magnitude with which a covariate is more associated with mortality vs. statistical temporary emigration. Positive (*blue*) values suggest a metric is more linearly associated with mortality than temporary emigration, while negative (*red)* values suggest the opposite.

#### 4.3.2 Data Leakage

Figure 5 shows a large change in the log-odds of dying vs changing closeness centrality, whereas the log-odds gradient over Δ-degree was much shallower and may just have been driven by a single data-point. This suggests that animals who died did not necessarily have a reduced number of total interactions and survey-encounters prior to death. Had this not been the case, a strong functional relationship between log-odds-death and degree may have revealed problems with data leakage. This suggests that social processes were changing prior to death, and metrics were not merely responding to a reduction in the underlying number of survey-encounters reduced observed-associates.

#### 4.3.3 Age

Age is an obvious predictor of mortality; it is interesting only inasmuch as it interacts with social metrics. For instance, there was a relatively smooth interaction between age and time-spent-only (Δ-p(alone), Figure) that may actually be indicative of the same fundamental aging process (older animals are naturally going to have fewer companions as they are lost to old age), whereas Δ-closeness centrality (Figure) and the widowhood-effect metric (Figure) showed more consistent effects regardless of age.

## 5 Discussion

Among large-brained social animals, populations of bottlenose dolphin serve as model systems for studying sociality. Within population, they exhibit high-levels of social-heterogeneity, which offers fruitful variation to explore connections between social phenotypes and individual life-history outcomes; for example, heterogeneity by sex, foraging groups, and high-levels of individual variation [Smolker et al., 1992a, Connor et al., 1992, Mann et al., 2000, Gibson and Mann, 2008, Connor and Krützen, 2015]. The question naturally arises how such within-population heterogeneity is maintained, especially through the lens of fitness outcomes and selection pathways. Natural selection for this kind of relational-complexity may have piggy-backed on the evolution of other cognitively-demanding traits and their respective fitness-benefits (e.g., problem-solving) [Jolly, 1966, Borrego and Gaines, 2016, Benson-Amram et al., 2016]. Or, it could be that complex sociality and social structure have their own fitness benefits and selection pathways [Brent et al., 2013, Gilby et al., 2013, Fowler et al., 2009, Brent, 2015], such socially-mediated access to mating opportunities or as advantaged survival through “social buffering”[Wittig et al., 2016, Silk et al., 2010].

In this study, we offer support for the latter, with an interest in how sociality changes over time, and how such changes are predictive of mortality (a fitness-proxy). Specifically, we offer support for a type of “widowhood effect” in males, as well as negative effects from declines in closeness centrality. We build upon a wide basis of research that previously focused on proving the existence of “non-random” associations, and ultimately moved towards hypothesis-driven inquiries into the evolution, mechanisms, and functions of social structure and social connectivity [Connor, 2007, Borrego and Gaines, 2016, Brent, 2015, Benson-Amram et al., 2016].

Most of the existing research on wild populations have focused on static measures of social bonds and social structure. Partially, this may be due to some aspects of sociality being relatively stable over life-times, such as differences between sexes or dominance-hierarchies [Silk et al., 2010]. Furthermore, popular lower-order measures of sociality (e.g., individual-traits like gregariousness, number of bonds, bond-strength, etc.) may be more stable in time, as they may reflect individuals’ heritable variation, like the non-human analogues of psychometric features [Jang et al., 1996]. However, certain aspects of social variation, such as the higher-order features that depend on indirect connections within a network, should be less stable over a lifetime. Such dynamics could be due to individual maturation, aging and reproduction status [Turner et al., 2017], while certain changes may be more reflective of complex social processes. For example, changes in social status and its consequences on depressed physiological states [Creel et al., 2013, Knight and Mehta, 2017]. Such longer-term dynamics in higher-order measures of sociality are difficult to quantify with short-term sparse data-sets of fission-fusion populations. Importantly, statistical inquiries into such social processes have a fundamental challenge to do with the imprecision of available social metrics and the inability to identify causal directions between changes in sociality and fitness-outcomes.

Given these challenges, we employed two machine-learning statistical frameworks to build evidence about the relationship between changes in higher-order social processes and individuals’ mortality status. Both inferential frameworks are commonly used in behavioral ecology, but we emphasize how they provide different interpretations of quantitative results. In the following discussion, we also use these two frameworks to suggest multiple competing interpretations of our quantitative results, especially when trying to account for the imperfect nature of social-metrics vs. underlying social processes. Behavioral ecologists are increasingly having to reckon with the tenuous connection between social-processes and social-metrics-as-hypotheses, and it is important to take stock of how this deficiency manifests in conventional hypothesis-testing frameworks [Brent et al., 2013]. For example, interpretations of the quantitative results of closeness centrality must contend with the fact that the metric lacks a straight-forward interpretation [Brandes et al., 2016] and may be redundant to other non-target measures [Rankin et al., 2016, Firth Josh A. et al., 2017].

The conservative interpretation of our quantitative results is that all we can say is that there is some undifferentiated importance to changes in sociality, while not necessarily being able to ascribe evidence for or against any specific metric/process ^6^ (i.e., we can only predict outcomes, not refute or confirm hypotheses). This was operationalized through our predictivist framework: a prediction-optimized “black-box” style of algorithmic inference, that merely provides an overall proof-of-concept that dynamic higher-order social metrics have predictive power on dolphin mortality. This is scientifically less satisfactory than inferential frameworks that focus on specific hypotheses, but also relies on less assumptions, and is therefore more robust in its limited claims on “truth”.

The more liberal interpretation is operationalized through our stability-selection inclusion probabilities, which serve as (approximate) confirmation functions [Hawthorne, 2011] for specific metrics-as-hypotheses. In the following section, we discuss how some specific conclusions are justifiable in the face of statistical difficulties that are rife in social network analyses, such as metric non-specificity and multi-collinearity.

### 5.1 Widowhood effect

Our metric for the (putative) widowhood effect had very high support and was most prevalent in males (as revealed in marginal plots, Figure 4). To understand why the metric was so predictive of individuals’ mortality, we offer three interpretations: the literal interpretation; a confounding “third-variable problem” via social clustering; and a more general erosion of social status.

Our favored interpretation, the widowhood effect, posits that the loss of alliance-members, especially among males, elicits a negative long-term physiological response and causes physical vulnerabilities. It could be that widowed males are more vulnerable to attacks by other males and predators, and/or that the loss of associates caused detrimental physiological stress, such as the loss of status and connection to the broader network. The crux of the effect is that dolphins have unique social identities and it is the loss of *specific* relationships that is detrimental. This is distinct from a more general deterioration of one’s social environment, such as reductions in group size, reduced time spent with others (strength), loss of the number of associates (degree), turnover in companions (social variance), and other low-level measures of sociality and undifferentiated relationships. In other words, individual dolphins are not substitutes for one another. This contrasts with societies of high-organizational complexity [Lukas and Clutton-Brock, 2018], where there are no unique fitness benefits from specific social bonds among undifferentiated individuals. This is a key distinction from our alternative explanations (below) that have no underpinning in individual identities nor relational complexity (individually-differentiated, long-term bonds).

Our second explanation for the (putative) widowhood-effect metric could be due to confounding by a third variable: the same statistical result could arise if a third-variable influenced both the metric and mortality (known in psychology literature as the “third variable problem”: TVP). For example, TVP could arise if any deleterious physical phenomenon clusters by social group, such as local resource stress, contagious disease, shark attacks. Social-clustering of a TVP could lead to a strong statistical association between a metric and mortality, even if there was no functional connection between changes-in-sociality and mortality; rather, deaths just cluster by social groups. If TVP were a significant phenomenon, we would expect that factors such as age and time-as-a-covariate would correlate with our measure of the widowhood effect, and would likewise have high predictive power (which was somewhat demonstrated in the results). However, we think TVP was less important as an explanation, according to the following reasoning: the existence of social-clustering TVP should be more acute among females, not males, given how the former are more specialized in their foraging behavior, more insular by geography and matrilines, and less socially promiscuous than males [Smolker et al., 1992a, Connor et al., 1992, Mann et al., 2000, Gibson and Mann, 2008, Connor et al., 2011]. In other words, third-variable exogenous shocks are more likely to be socially and locally clustered among females. However, our widowhood effect metric was more pronounced among males, which leads us to somewhat downplay the social-clustering TVP as a principal driver the widowhood-effect result.

Our third competing explanation was that the appearance of the widowhood-effect was not necessarily due to any specific identifiable social process; instead, our metric was just correlated with other unspecified social processes, such as changes in strength, degree, time-spent-alone, etc. In this case, poor specificity results in broad sensitivity to less-interesting and undifferentiated forms of sociality change. If the metric is just confounded with changes in low-level individual-attributes like strength and degree, it is less interesting for hypothesis-focused research trying to advance understanding about the fitness-consequences of sociality and the heritability of higher-order social processes [such as centrality and indirect-connectedness McDonald, 2007, Gilby et al., 2013, Brent et al., 2013]. The results from our RVI interactions suggest that this interpretation has some merit. For example, the widowhood-effect metric had a strong interaction with changes in the amount of time that an individual spent alone (Δ-p(alone)).

However, the results from the dropout-RVI statistic reveas that the widowhood-effect metric *did* have a unique contribution towards predictive performance, and which could not be replaced by *many* non-linear combinations of other covariates. This independent contribution was strong even though there was high statistical multi-collinearity and converns about the the metric’s specificity and operationalization. Therefore, we suggest the our literal interpretation is most compelling, and that the relative *un*importance of other lower-level metrics suggest that the mundane interpretations (such as TVP) are not very convincing as explanations.

### 5.2 Closeness Centrality

Centrality metrics are prominent features in studies about sociality and fitness-surrogates, such as betweenness centrality, information centrality, and eigenvector centrality [McDonald, 2007, Stanton and Mann, 2012, Gilby et al., 2013]. These are exciting metrics because, unlike lower-order metrics which pertain to single dimensions (e.g., more or less bonds, weaker or stronger bonds, larger or smaller group-size) network-position metrics reveal the ways in which seemingly low-degree individuals can occupy “central” positions, such as being brokers of information (i.e, betweenness) or dependent upon intermediaries [e.g., closeness Brandes et al., 2016]. Such metrics should also be less stable in time, given their complex dependence on communities of individuals within networks; this instability offers an interesting natural experiment to look for statistical correlates to fitness-surrogates. We therefore expected centrality metrics to be highly significant predictors. However, we were surprised by the very strong evidential support for change-in-closeness centrality (the sex-homophily version). Unlike betweenness or eigenvector centrality, which have straight-forward interpretations, the interpretation of closeness centrality is less obvious [Brandes et al., 2016, Firth Josh A. et al., 2017]. The naive interpretation is that as a dolphin becomes more dependent on intermediaries for connections (low centrality), and is more socially-distant from all other individuals in the network, he is at a higher risk of dying (the alternative direction of causality should also be considered, as we do below). This interpretation is not implausible, given how dolphins devote a lot of energy socializing and maintaining relationships: it could be that when one’s social position erodes (as tracked by centrality), a dolphin is less useful, has less social status, and has more concentrated reliance on fewer individuals. We interpret this as a loss of social status, which can have harmful hormonal and physiological consequences in others species[Sapolsky, 1982]. Previously, we wrote skeptically about the utility of the closeness centrality metric [Rankin et al., 2016], claiming it was redundant to low-level non-network measures. Therefore, we are cautious in placing too much emphasis on the above naive interpretation of closeness-centrality. Instead, we suggest it is a good general and concise univariate measure for the entire social network and a node’s position within it. In other words, the metric integrates low-level social information like strength and degree [Rankin et al., 2016] as well as indirect connectedness [Firth Josh A. et al., 2017]. Therefore, measuring a change in closeness centrality could be a measure of a more general disruption of one’s social environment, including both low-level dyadic relationships as well as higher-order network position. We think it is more likely that the direction of causality is in the opposite direction, whereby closeness centrality could be merely tracking an animal’s social erosion as their health deteriorates into death; for example, other animals may avoid terminally-ill individuals [see Miketa et al., 2017, for one notable account of such an occurrence].

Nonetheless, despite the correlation that closeness centrality has with other metrics, the quantitative results strongly support the notion that closeness has some unique explanatory power and predictive ability, beyond all other non-linear combinations of strength, degree, eigenvector centrality, reach, betweenness, etc. We made a further effort to try and explain the metric by correlating it to other social measures in a more fine-grained social dataset[Galezo et al., 2018], as well as other metrics of social disruption. However, we could not find a strong and convincing reductivist explanation. Therefore, we merely accept that the metric is of interest from a predictivist stand-point, and suggest cautiously that it may be deserving of further fine-scale scrutiny. In a purely predictivist framework, it seems to have very high importance and should be investigated in studies of sociality and fitness-surrogates.

### 5.3 Metric Imprecision

The issues of multi-collinearity and low-specificity are pervasive in observational studies of animal sociality. From a predictivist perspective, neither of these issues are relevant if our principal primary goal to predict individual outcomes. Such “black-box” use-cases are common in applied machine-learning and can be useful for conservation managers or population modelers. For example, conservationists may value probabilistic assessments about which individuals are likely to die, or modelers may like to augment population-biology models (such as capture-recapture models) with socially-derived information about mortality status. In this way, applied sociality research can augment the decision making of conservation managers [Snijders et al., 2017], irrespective of our misgivings about metrics’ specificity and collinearity.

However, for hypothesis-focused research based on observational data, having low specificity of metrics and statistical multi-collinearity erodes our power to discriminate among competing hypotheses. This is especially corrosive to the falsificationist scientific framework and the method to advance science through refuting hypotheses. For example, consider our hypothesis about the importance of daughters for mothers’ survival: we hypothesized that surviving daughters serve as social anchors to the broader female community. However, we found no evidence for a marginal effect of these metrics, given our 3 different operationalizations and independent investigations of each metric. The metrics provided little-to-no predictive power, and our inclusion probabilities suggested that there was no basis to believe in their importance.

Does such quantitative evidence refute our hypothesis? There always exists the fundamental insecurity about the sufficiency of such metrics to actually measure their intended social process, and thus refuting them only refutes the metrics, not the process. This type of insecurity is not unique to our methods, and is widespread in the biological sciences [e.g., Pearce, 1990, argues that biological hypotheses are rarely falsifiable, and this has forced biologists to abandon the Popperian falsification framework of science in favor of the much-maligned NHST framework], but the issue is especially problematic for social network metrics with low specificity and high multi-collinearity.

At least on the confirmatory side^7^, we used a few different strategies to help build a convincing argument for the importance of higher-order sociality, such as the widowhood effect. Specifically, by including two inferential frameworks, one which was conservative and only emphasized the predictive importance of changes in sociality, and the second framework being more ambitious in trying to ascribe probability-values to metrics-as-hypotheses. We also investigated the role of confounding processes and artefacts, such as statistical temporary migration, problems due to unknown mortality-states, data-leakage, and age. Ultimately, these kinds of hypotheses may be at the limits of what can be robustly evaluated using sparse observational data. Future analyzes will need to leverage finer-scale data and develop precise and specific social metrics.

## Footnotes

1 One commonly-raised concern about overlapping windows is whether it violates assumptions of statistical independence among observations (in contrast to previous studies which used three-year non-overlapping windows). We believe this concern is unfounded. Rather, the issue of overlapping versus non-overlapping windows reflects a trade-off between bias versus variance in the relationship between measured social-changes and point-in-time death events. Non-overlapping windows create an arbitrary bisection of an animals time-series and life-history (such as reproductive events or social changes) as well as increasing the temporal-distance between relevant social-changes and a death event. Such bias dampens the statistical power to correlate death-events and life-history changes and/or sociality changes.

2 Ecologists more familiar with the AIC should note that the Expected Risk is the same target that the AIC seeks to approximate and minimize [Eqn. 1.1; Akaike, 1998], whereas, for boosting, the target 𝔼 [*−*log(*ℒ*)] can be approximated with an 100-fold bootstrap

3 We adopted an approximate Bayesian interpretation of stability selection inclusion probabilitiesDraper [2010] because we have a Bayesian philosophy about the nature of probabilities and hypotheses. We consider inclusion probabilities to quantify our degree-of-belief about *specific* hypotheses from a *specific* study-system. This is a fundamentally Bayesian world-view, despite the underlying technology being based on frequency resampling. Although approximate, the framework contrasts sharply with the Neyman-Pearson notion of science being about long-run ‘error control’ We think the error-control interpretation of stability selection has merit in large-scale repeatable industrial applications or quality-control programs (as in the original paper about bacterial gene-expression and riboflavin production), but less so for inference about specific hypotheses in a single population of dolphins.

4 The number of false-positives and false-negatives goes to zero with probability approaching 1 as *n* gets large.

5 given assumptions of linearity and exchangeablility of base-learners

6 By ‘not robust’, we mean that our predictivist framework lacks the rigorous quantification of “significance” or “consistency” or analogous theoretical guarantees on Type-I errors, etc., which are important for hypothesis-driven research

7 Here, “confirmation” refers to the Bayesian notion of being rational agents who bring one’s degree-of-belief into alignment with one’s posterior inference. Such subjective beliefs are not static and are updated via posterior inference Hawthorne [2005, 2011].

